# Sex-specific metabolic regulation by the Drosophila RNA-binding protein Nab2

**DOI:** 10.64898/2026.09.25.754360

**Authors:** Jordan N. Goldy, Heath Dunlop, Dev D. Patel, Rebecca M. McSweeney, Sara W. Leung, David U. Gorkin, Anita H. Corbett, Kenneth H. Moberg

**Affiliations:** Department of Biology, Emory College of Arts and Sciences, Atlanta, GA 30322; Department of Cell Biology, Emory University School of Medicine, Atlanta, GA 30322; Graduate Program in Biochemistry, Cell and Developmental Biology, Emory University, Atlanta, GA 30322; Graduate Program in Genetics and Molecular Biology, Emory University, Atlanta, GA 30322

## Abstract

Conserved RNA binding proteins (RBPs) regulate key steps of gene expression including mRNA processing, export, localization, stability and translation. Human ZC3H14 is a conserved RBP that regulates pre-mRNA processing in neurons and loss of ZC3H14 leads to neurological defects. Studies of Nab2, the *Drosophila* orthologue of ZC3H14, have identified potential target RNAs involved in metabolism, suggesting Nab2 may influence neurometabolic circuitry. Here, we show a female-specific increase in *dilp2* and *dilp5* mRNA levels. The dilps encode insulin-like peptides that signal from the brain insulin producing cells (IPCs) to peripheral tissues. *Nab2^null^* females have enlarged lipid droplets in the fat body, a tissue analogous to human adipose tissue and liver. Notably, neuronal depletion of Nab2 increases lipid droplet size while neuronal expression of Nab2 in *Nab2^null^* female rescues this phenotype supporting a role for Nab2 in a neuronal circuit that regulates *dilp* levels. Furthermore, depletion of *dilp2* or *dilp5* from IPCs rescues the enlarged lipid droplet phenotype in *Nab2^null^* females indicating that elevated *dilp2*/*dilp5* contributes to enlarged lipid droplets. Together, these data support a female-specific role for Nab2 in brain neurons to support insulin signaling and fat storage, expanding the known functions of RBPs linking neuronal function and metabolic homeostasis.

**Significance Statement:**

- *Drosophila* Nab2, orthologue of the human RBP ZC3H14, which is lost in an inherited form of intellectual disability, regulates a specific subset of neuronal RNAs.
- This study identifies a female-specific role for Nab2 in regulating systemic metabolism by maintaining insulin-like peptides *dilp2* and *dilp5* mRNA levels in the brain and lipid homeostasis in the fat body, a tissue analogous to human adipose tissue and liver.
- These findings suggest that Nab2 has a role in a female-specific circuit that helps to maintain metabolic homeostasis involving the insulin signaling pathway and lipid storage in the fat body.

## Introduction

Ubiquitously expressed, evolutionarily conserved RNA-binding proteins (RBPs) play essential roles in regulating gene expression, guiding all steps in mRNA processing including splicing, polyadenylation, export, localization, translation and stability (Glisovic et al. 2008, Corbett 2018, He et al. 2023, Lancaster et al. 2024). Critically, RBPs regulate spatiotemporal patterns of gene expression to ensure proper cellular homeostasis and guide tissue-specific functions, making them particularly important for brain development. (Kishore et al. 2010, Darnell 2013, Conlon et al. 2017, Corbett 2018, He et al. 2023). As a result, mutations in genes encoding RBPs often lead to neurodevelopmental disorders (Cooper et al. 2009, Pak et al. 2011, Bardoni et al. 2012, Darnell 2013). A growing body of evidence also shows that the brain can regulate signaling events in various other tissues for proper function and survival (Morton et al. 2006, Ahima et al. 2008, Zhou et al. 2023, Zou et al. 2024). More specifically, mutations in genes encoding RBPs with roles in neurodevelopment have also been linked to systemic metabolic defects (Benegiamo et al. 2018, Leboucher et al. 2019, Ceron-Codorniu et al. 2024). RBPs such as FMRP (Hagerman et al. 2017), TDP-43 (Jo et al. 2020), and NONO (Mircsof et al. 2015) that are linked to neurological disorders have also been implicated in metabolic regulation involving insulin signaling and lipid metabolism (Benegiamo et al. 2018, Leboucher et al. 2019, Ceron-Codorniu et al. 2024). Further strengthening the connection between RBPs and metabolism, a recent large-scale exome sequencing study linked variants of a protein required for post-transcriptional regulation to sex-specific fat accumulation in females but not males (Kaisinger et al. 2023). These findings highlight a gap in our understanding of how loss of a single RBP with important functions in neurons, can lead to specific metabolic dysfunction in distant tissues and how these functions maybe be inter-related or sex-specific.

One RBP that has documented roles in neurons is the Zinc finger Cys-Cys-Cys-His-type containing 14 (ZC3H14; also termed MSUT2) (Leung et al. 2009, Kelly et al. 2010, Wheeler et al. 2019, Shelby et al. 2026). Mutations in the gene encoding ZC3H14 are associated with a neurodevelopmental disorder in humans (Pak et al. 2011, Kelly et al. 2012), illustrating the importance of the RBP in the central nervous system. ZC3H14 is an evolutionarily conserved RBP with homologs present in *Mus musculus* (Zc3h14) (Guthrie et al. 2011, Pak et al. 2011, Soucek et al. 2016), *Caenorhabditis elegans* (SUT-2) (Guthrie et al. 2009, Fasken et al. 2019), *Saccharomyces cerevisiae* (Nab2) (Fasken et al. 2019), *Saccharomyces pombe* (Nab2) (Grenier St-Sauveur et al. 2013), and *Drosophila melanogaster* (Nab2) (Pak et al. 2011, Kelly et al. 2016, Fasken et al. 2019, Lee et al. 2020, Corgiat et al. 2021, Corgiat et al. 2022, Rounds et al. 2022, Jalloh and Lancaster et al. 2023, Lancaster et al. 2024). Research using various model systems has provided insight into molecular functions of ZC3H14 including regulation of poly(A) tail length (Pak et al. 2011, Kelly et al. 2014, Gabs et al. 2025), alternative splicing (Jalloh and Lancaster et al. 2023), control of m^6^A levels (Jalloh and Lancaster et al. 2023), co-transcriptional RNA processing (Alpert et al. 2020) and compaction/packaging of mature transcripts for export (Aibara et al. 2017). Together, these findings highlight the conserved role for ZC3H14/Nab2 in RNA metabolism and contribute to our understanding of how loss of an RBP may underlie neurodevelopmental disorders. The connection of various RBPs to metabolic defects, raises the question of whether ZC3H14/Nab2 loss may also cause metabolic dysfunction due to its important role in regulating a variety of RNA processing events.

*Drosophila* provides a genetic tractable model for investigating the role of ZC3H14/Nab2 as a regulator of target RNAs. Our previously developed *Drosophila* model that lacks *Nab2* mRNA and protein (*Nab2^null^*) provides a model of ZC3H14 loss of function associated with human neurodevelopmental disease (Pak et al. 2011). Studies in the fly model revealed that loss of Nab2 results in impaired neuronal function (Pak et al. 2011, Corgiat et al. 2022, Lancaster et al. 2024), a decrease in viability (Pak et al. 2011), and changes in the brain transcriptome including defects in splicing of ∼150 mRNAs (Jalloh and Lancaster et al. 2023). Importantly, human ZC3H14 can functionally substitute for *Drosophila* Nab2 in neurons and rescue many of the phenotypic defects observed in the *Nab2^null^* flies (Kelly et al. 2014). These data are consistent with an evolutionarily conserved role for ZC3H14/Nab2 in many aspects of RNA processing within neurons. Despite the work investigating the role of Nab2 in neurons (Pak et al. 2011, Kelly et al. 2012, Kelly et al. 2016, Corgiat et al. 2021, Rounds et al. 2022, Jalloh and Lancaster et al. 2023, Lancaster et al. 2024), a potential role for Nab2 in metabolic homeostasis has not been explored. Interestingly, RNA sequencing of *Nab2^null^* fly heads compared to Control heads revealed that loss of Nab2 affects multiple metabolic transcripts including key regulators of lipid metabolism and insulin signaling (Jalloh and Lancaster et al. 2023). Exploiting the established *Drosophila* Nab2 model has the potential to provide insight into human ZC3H14 as a metabolic regulator as many metabolic processes are conserved between flies and humans (Chatterjee et al. 2021, Kim et al. 2021, Cesur et al. 2023, Moon et al. 2026). Among these conserved metabolic pathways are insulin signaling, lipid storage, and carbohydrate metabolism further strengthening rationale to explore Nab2-mediated regulation of metabolic processes (Yamaguchi et al. 1995, Garofalo 2002, Graham et al. 2017, Viola et al. 2023).

Insulin signaling is well conserved between *Drosophila* and humans with the insulin producing cells (IPCs) in the *Drosophila* brain playing a similar role to the pancreatic β islet cells in humans (Kréneisz et al. 2010). The IPCs produce and secrete *Drosophila* insulin like peptides (Dilps) to regulate lipid storage in distant tissues (Géminard et al. 2009). The main lipid storage tissue in flies is called the fat body, which is analogous to human adipose tissue and liver (Arrese et al. 2010). Binding of the Dilps to the insulin-like receptor on the surface of target cells activates a conserved phosphorylation cascade involving phosphoinositide 3-kinase (PI3K) and Akt (Garofalo 2002). This signaling pathway promotes nutrient uptake and lipogenesis while inhibiting lipolysis leading to increased lipid storage (Teleman 2009, Biswas et al. 2025). In *Drosophila*, Dilp2, Dilp3 and Dilp5 are produced and secreted from the IPCs to regulate lipid storage in the fat body (Géminard et al. 2009, Viola et al. 2023). Dilp2 is one of most abundant and well-studied of the Dilps with expression throughout larval development and adulthood to regulate body size, lifespan and lipid metabolism (Géminard et al. 2009, Kannan et al. 2013, Semaniuk et al. 2021). Dilp3 contributes to metabolic homeostasis and can compensate for reduction in the expression of Dilp2 or Dilp5 signaling during specific nutritional and genetic conditions (Broughton et al. 2008, Kim et al. 2015). Dilp5 promotes larval growth through nutrient-dependent growth responses and contributes to lipid and carbohydrate metabolism (Géminard et al. 2009, Nässel et al. 2013, Post et al. 2018). While all three of these Dilps are expressed in the IPCs (Brogiolo et al. 2001), Dilp2 and Dilp5 are considered the primary regulators of nutrient-dependent growth and metabolic homeostasis (Post et al. 2018).

In both humans and *Drosophila*, females and males generally maintain different levels of lipid storage for energy and the sexes have distinct insulin-signaling responses (Power et al. 2008, Wat et al. 2020, Diaz et al. 2023). This difference can be highlighted by the expression of sex-specific hormones, gene expression and reproductive demands that modulate energy utilization and storage (Power et al. 2008, Mauvais-Jarvis 2015). As mentioned earlier, previous RNA sequencing revealed an increase upon loss of Nab2 in adult female fly heads of the steady state levels of mRNAs involved in lipid metabolism such as *Drosophila* insulin-like peptides 2 and 5 (*dilp2* and *dilp5*) (Jalloh and Lancaster et al. 2023). Therefore, we can leverage the *Drosophila* model to investigate the role that ZC3H14/Nab2 may have in regulating insulin signaling and lipid metabolism..

Here, we employ both genetic and molecular techniques to assess the link between Nab2 and *dilp2/5* mRNA in the brain and lipid storage in the fat body. First, we find that these transcripts are only elevated in female adult *Nab2^null^* fly heads and not in *Nab2^null^* male heads. We establish that these changes in *dilp* transcript levels in females are not specific to adulthood but also occur at an earlier developmental time point. We find that female *Nab2^null^* fat bodies show an increase in lipid droplet size compared to Control that is not observed for male fat bodies. Intriguingly, neuronal depletion of Nab2 increases lipid droplet size in the fat body but depletion from IPCs does not. Moreover, neuronal expression of a *Nab2* transgene in *Nab2^null^* larvae rescues the lipid droplet size defect in the female *Nab2^null^* fat body. Finally, depletion of *dilp2* or *dilp5* from the IPCs of a female *Nab2^null^* larval brain rescues the increase in lipid droplet size seen in the female *Nab2^null^* larval fat body, providing a causal link between elevated *dilp* levels and fat body lipid storage. Taken together, these data identify a sex-specific role for Nab2 in brain neurons that regulates systemic fat storage. Our results suggest that this sex-specific regulation of lipid metabolism may stem from a role for Nab2 in neurons that synapse on IPCs, or that regulate *dilp2* and *dilp5* transcript levels. These data support a role Nab2 as an RBP with important roles in both proper neuronal function and lipid metabolism.

## Results

### Nab2 regulates dilp2 and dilp5 steady state levels in female adult and larval brain

Gene Ontology (GO) analysis of previously acquired high-throughput RNA sequencing (RNA-seq) data from the heads of adult female and male *Nab2^null^* mutants and isogenic Controls (Jalloh and Lancaster et al. 2023) was performed for GO Biological Process terms to identify biological processes significantly overrepresented among the differentially expressed genes. This GO analysis revealed extensive changes in the fraction of the brain transcriptome that encodes metabolic regulators and enzymes (Figure 1A). To further focus this analysis on cellular energy balance and nutrient sensing pathways, *Nab2^null^* and isogenic Control head transcriptomes were analyzed for changes in GO terms related to lipid-associated biological processes (Figure 1B). This GO analysis revealed a significant enrichment for factors involved in cellular lipid metabolism and fatty acid metabolic processes among Nab2-regulated brain mRNAs relative to Controls, suggesting that Nab2 loss impacts lipid and energy metabolism. Interestingly, the enrichment for GO terms in lipid-related processes in *Nab2^null^* heads vs. Control heads is significantly greater in female heads as compared to male heads. Among the transcripts that showed statistically significant changes in steady-state levels within this dataset are two metabolic transcripts, *dilp2* and *dilp5 (*Jalloh and Lancaster et al. 2023). These mRNAs encode *Drosophila* insulin-like peptides (Dilps) that regulate lipid storage in the *Drosophila* fat body (Géminard et al. 2009, Viola et al. 2023), the primary metabolic and energy storage tissue in *Drosophila* (Arrese et al. 2010).

**FIGURE 1.**
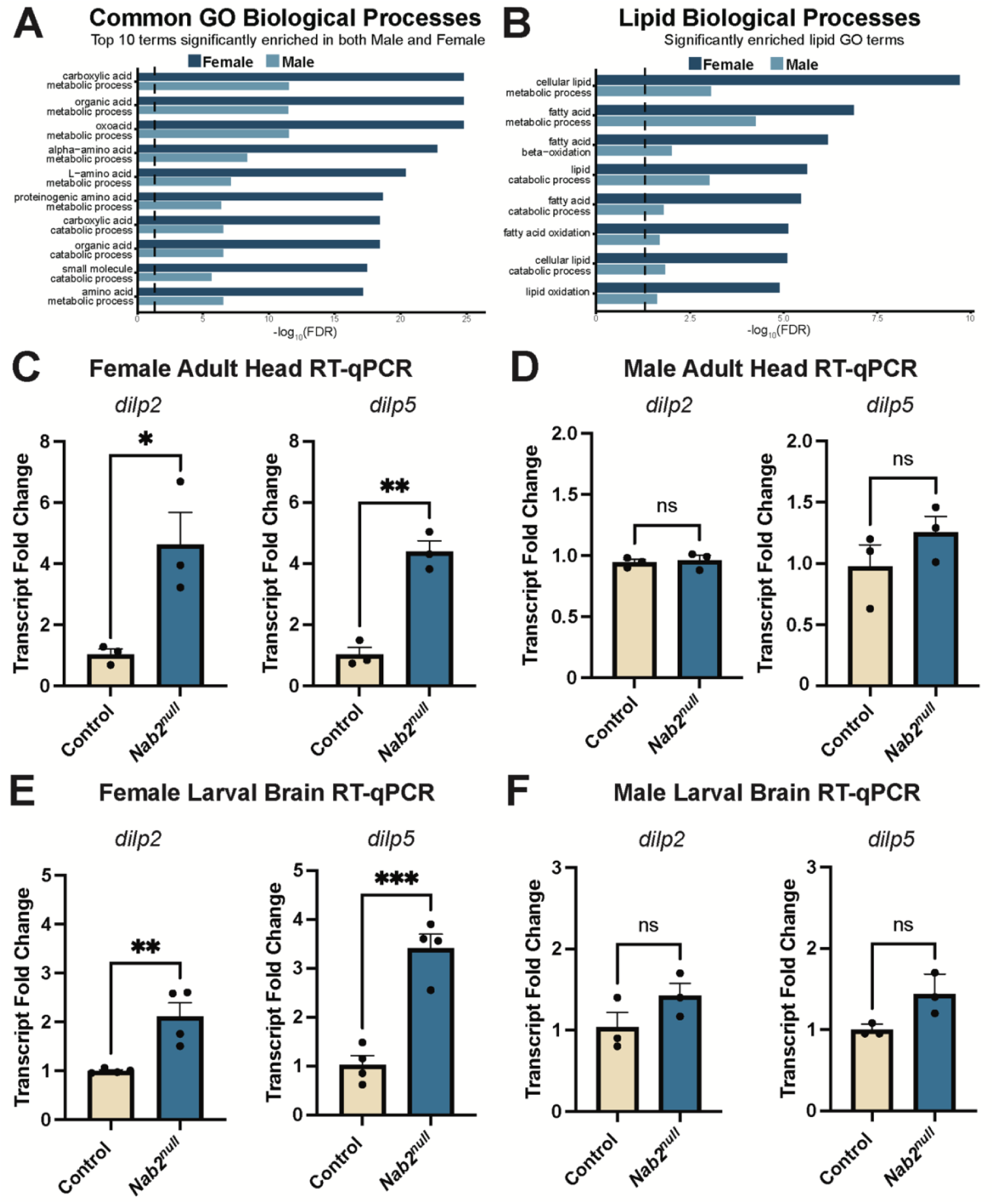
Nab2 regulates dilp2 and dilp5 transcript steady state levels in female adult and larval brains. GO enrichment was assessed for Biological Processes (A) and Lipid Biological Processes (B) to identify the significantly enriched terms (*p*-value (FDR) < 0.05) among the differentially expressed genes in Nab2^null^ females and males relative to sex-matched Controls. Bar plots display the -log_10_ transformed FDR values. Dashed line represents the significance threshold of 0.05. (C,D) qRT-PCR analysis analyzing the levels of *dilp2* and *dilp5* in Control and *Nab2^null^* (C) female and (D) male adult heads. qRT-PCR analysis was performed to analyze transcript levels of *dilp2* and *dilp5* in Control and *Nab2^null^* (E) female and (F) male larval brains. RNA from 20 adult heads and 25 larval brains was extracted from each genotype for each biological replicate. The Control was set to 1.0 in each case and results are plotted as Transcript Fold Change. A minimum of three biological replicates was performed for each experiment. Larvae were aged matched. Welches t-test was performed and significance values are indicated. (*, *p*-value <0.05) (**, *p*-value <0.005) (***, *p*-value <0.001). Error Bars represent s.e.m.

RT-qPCR analysis was used to confirm that *dilp2* and *dilp5* transcript levels are increased in adult female *Nab2^null^* heads relative to Control female heads (Figure 1C). In contrast, the steady-state level of these transcripts is not significantly altered in adult male heads (Figure 1D). As both *dilp2* and *dilp5* are expressed across developmental timepoints (Post et al. 2018), we also examined *dilp* transcript levels in the brain during the 3^rd^ instar larval phase. As shown in Figure 1E and 1F, this qRT-PCR analysis confirms that *dilp2* and *dilp5* steady-state transcript levels are elevated upon loss of Nab2 specifically in female larval brains and not in male larval brains. These data support a female-specific role for Nab2 in regulating *dilp2* and *dilp5* mRNA levels in the adult and larval brain.

### Loss of Nab2 enhances female larval size without affecting feeding behavior

Insulin signaling promotes fat storage in both humans and *Drosophila* (Teleman 2009) and varies across developmental stages and with altered feeding behavior (Sudhakar et al. 2020). To investigate potential changes resulting from altered lipid storage that could occur due to changes in levels of *dilp2* and *dilp5* transcripts, we measured overall weight and size of female and male 3^rd^ instar larvae, which corresponds to a stage of rapid growth (Church et al. 1966). Female *Nab2^null^* larvae show a statistically significant increase in overall larval weight compared to Control females (Figure 2A) while *Nab2^null^* and Control males show no statistically significant difference in weight (Figure 2B). This female-specific increase in weight is matched by an increase in *Nab2^null^* larval width in females with no corresponding statistically significant change in larval length (Figure 2C). No statistically significant differences in length or width were detected when Control and *Nab2^null^*males were compared (Figure 2D). As changes in *dilp2* and *dilp5* mRNA levels and larval weight and width could be a product of altered feeding behavior, we assessed food intake by rearing larvae on food containing Bromo-phenol Blue and then measuring absorbance of Bromo-phenol Blue in whole-animal lysates. Using this approach, we did not detect a significant difference in the amount of dyed food in the guts comparing *Nab2^null^* female (Figure 2E) or male (Figure 2F) *3^rd^* instar larvae relative to Controls. In sum, this analysis reveals that *Nab2^null^* female larvae show an increase in weight and width relative to Control that is not observed in males and that this increase in the size of the *Nab2^null^*female larvae is not due to increased food consumption and, thus, may be due to a role for Nab2 in regulating *dilp2/5* and lipid metabolism.

**FIGURE 2.**
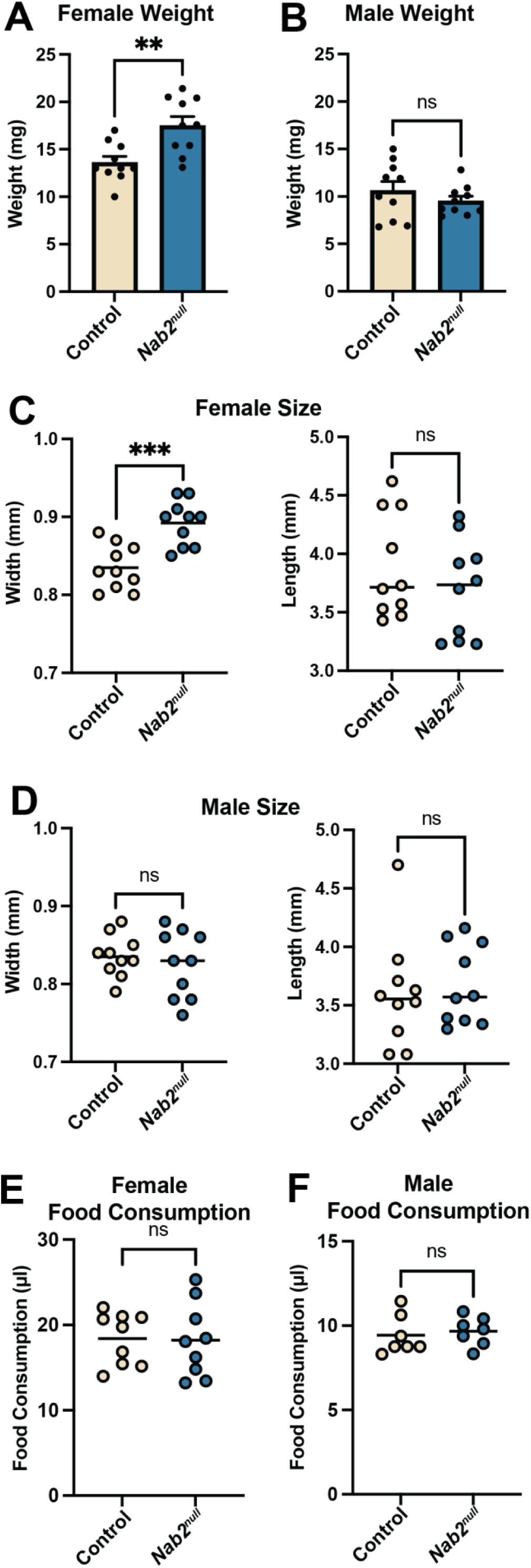
Loss of Nab2 enhances female larval size without affecting feeding behavior. Control and *Nab2^null^* third instar larvae, (A) female and (B) male, were weighed to measure overall larval weight. Each data point represents the weight of 10 larvae collected in an Eppendorf tube and weighed as described in Materials and Methods (10 biological replicates) (n=10). (C,D) Overall larval size was determined for Control and *Nab2^null^* third instar larvae by measuring the length and width of (C) females and (D) males (10 biological replicates). The amount of food in the guts of Control and *Nab2^null^* (E) female and (F) male third instar larvae was measured after the larvae consumed food dyed with Bromo-phenol Blue. Nine biological replicates for females and seven biological replicates for males were performed. Each experiment used aged match larvae. Welches t-test was performed and significance values are indicated. (**, *p*-value <0.005) (***, *p*-value <0.0003). Error Bars represent s.e.m.

### Average lipid droplet size in the female fat body is increased upon loss of Nab2

Dilp2 and Dilp5 peptides are produced and secreted from the brain IPCs into the hemolymph (Géminard et al. 2009). These peptides then bind the insulin-like receptor on the surface of specific cell types, including the fat cells in the fat body (Garofalo 2002). Increased insulin signaling in the fat body results in increased lipid storage in lipid droplets (DiAngelo et al. 2009), which are phase-separated droplets of neutral lipids surrounded by a thin layer of phospholipids and protein (Fujimoto et al. 2011, Kühnlein 2012). Given the increase in *dilp2* and *dilp5* transcript levels in *Nab2^null^* female brains, we examined overall lipid droplet size in dissected *Nab2^null^* and Control fat bodies using the neutral intracellular lipid stain Nile Red (Greenspan et al. 1985). Nile Red images for female (Figure 3A) and male (Figure 3B) fat body lipid droplets were acquired and analyzed using the ImageJ plugin, StarDist (Schmidt et al. 2018), which generates segmented images with exact measurements for each lipid droplet (Figure 3C). This analysis reveals that, lipid droplets in *Nab2^null^*female fat bodies are increased in average size compared to sex-matched Controls (Figure 3D), while there is no statistically significant change in lipid droplet size upon loss of Nab2 in males (Figure 3E). As triacylglycerides (TAGs) are the main neutral lipid found in the core of lipid droplets (Fujimoto et al. 2011), we measured levels of TAGs per unit protein in lysates of 3^rd^ instar larvae using an *in vitro* TAG assay (Beshel et al. 2017). This *in vitro* assay reveals an elevation in TAG concentration per unit protein in *Nab2^null^* female larvae but not males when compared to sex-specific Controls (Figure 3F). These data support a role for Nab2 in regulating fat body lipid droplet size and TAG levels in a sex-specific manner.

**FIGURE 3.**
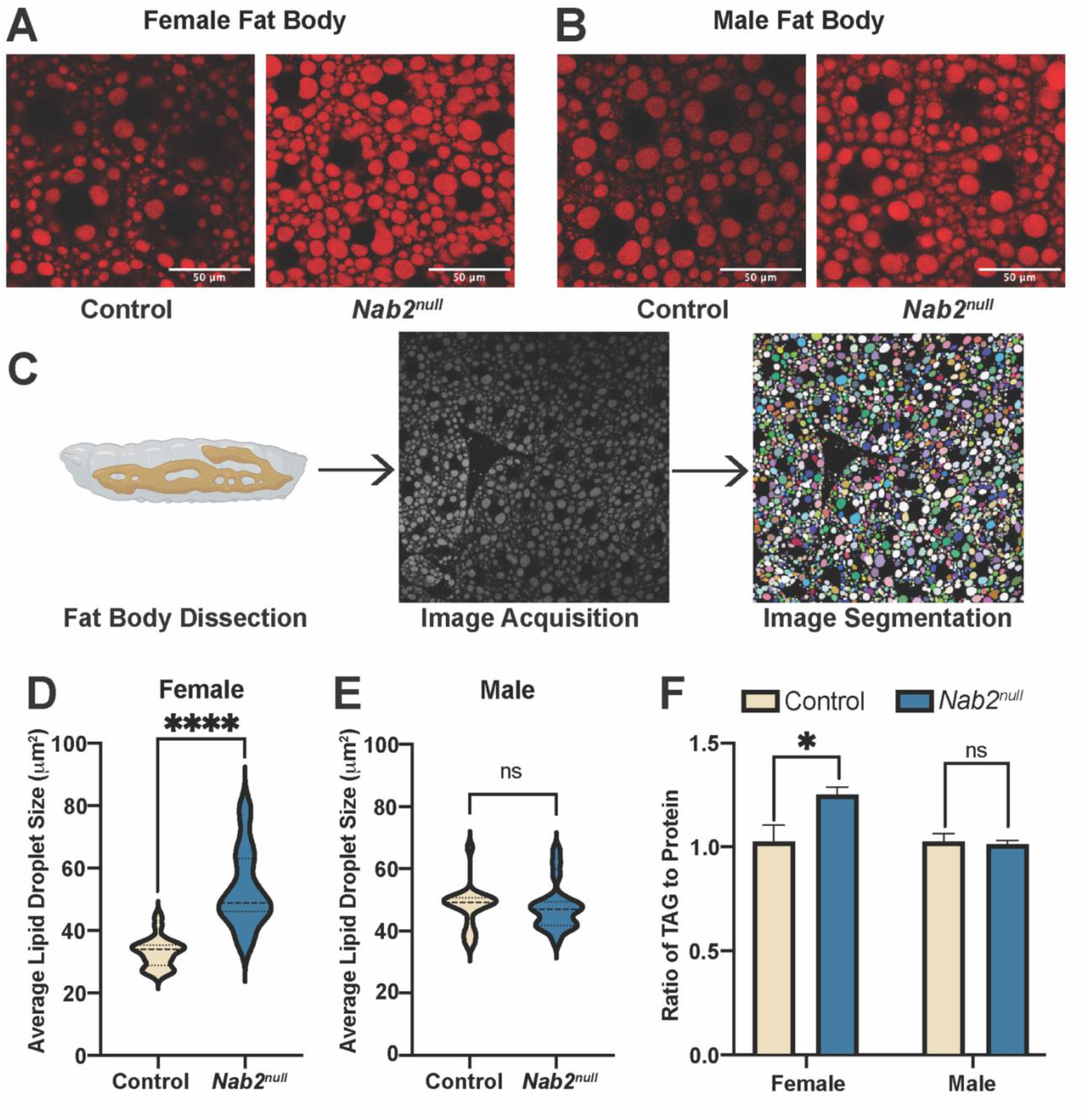
Average lipid droplet size in the female fat body is increased upon loss of Nab2. (A) Female and (B) male Nile Red-stained images for Control and *Nab2^null^* third instar larval fat bodies. Black circles represent nuclei of a single fat cell. (C) Depiction of lipid droplet size analysis. Third instar larval fat bodies were dissected (Created in BioRender. Goldy, J. (2026)), stained with Nile Red to visualize lipid droplets and images were segmented in Fiji using StarDist plugin to acquire the size (μm^2^) of each lipid droplet in an image. Average lipid droplet size for Control and *Nab2^null^* third instar larval fat body in (D) female and (E) male larvae. Each data point represents the average lipid droplet size for a single larval fat body. For this analysis, 20 age-matched biological replicates were used for each genotype for females and males. Welches t-test was performed and significance values are indicated. (****, *p*-value <0.0001). Error Bars represent s.e.m. and Scale bar = 50 μm

### Loss of Nab2 leads to lipid accumulation in hemolymph

A hallmark of obesity in humans is an increase in lipid storage and circulating lipids (Banerjee et al. 2025). This phenotype has also been observed in *Drosophila* where dysregulation of fat storage leads to an increase in circulating lipids in hemolymph (Walls et al. 2013). Due to the increased lipid storage in the fat body of the *Nab2^null^* female larvae, we tested whether the hemolymph from these flies also shows an increase in lipid content. For this analysis, hemolymph was collected from Control and *Nab2^null^* female larvae and stained with Nile Red to visualize lipid droplets comparing Control and *Nab2^null^* female larvae (Figure 4, A and B). As the Control sample appears to show a decrease in lipid droplets, we quantitated the results (Figure 4C) and found an increase in the amount and size of lipid droplets circulating in the *Nab2^null^* female hemolymph compared to Control. These data support a role for Nab2 in lipid homeostasis both in the fat body and in controlling lipid circulating in the hemolymph, both of which are markers for obesity in humans (Awari et al. 2025).

**FIGURE 4.**
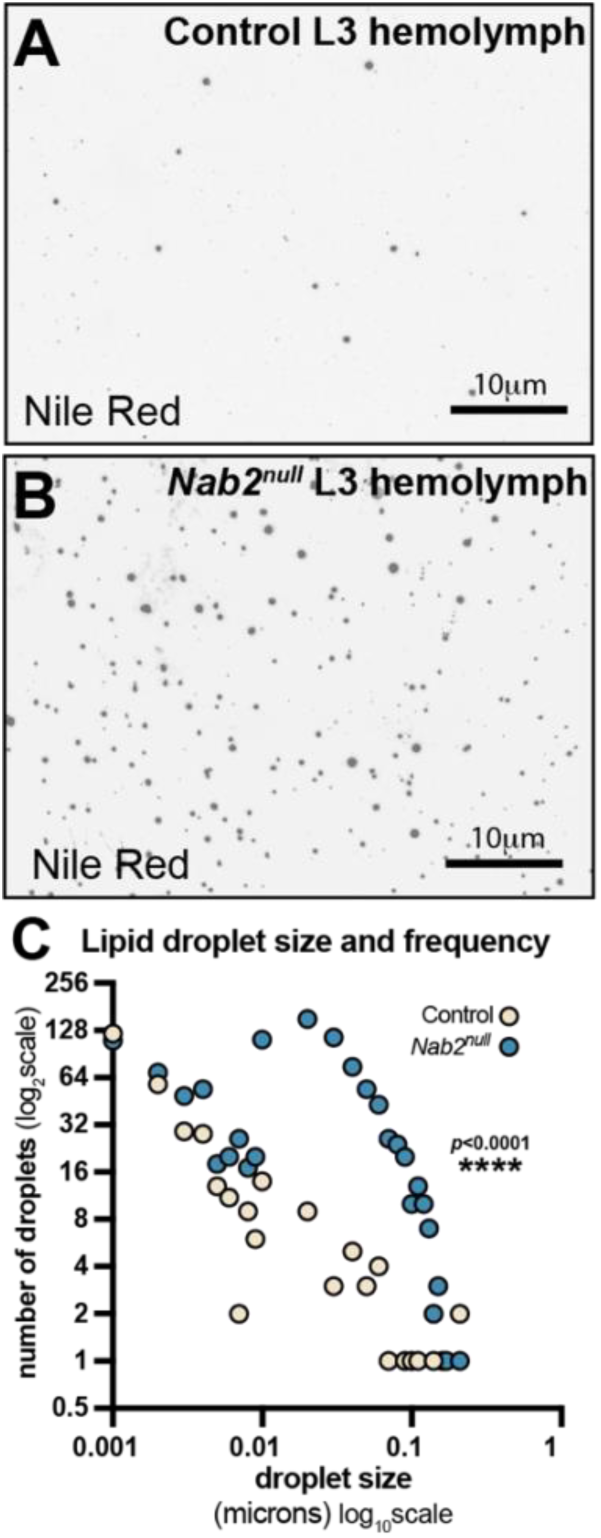
Loss of Nab2 leads to lipid accumulation in hemolymph. Hemolymph collected from (A) Control or (B) *Nab2^null^* 3^rd^ instar female larvae were collected and stained with the neutral intercellular lipid stain Nile Red. (C) Lipid droplet size was calculated for lipids detected in the hemolymph for Control and *Nab2^null^* female larvae. The plot shows the distribution of lipid droplet sizes for each genotype. Lipid droplets were grouped into size bins along the x-axis and the y-axis represents the number of lipid droplets detected within each size bin. Data are shown for three biological replicates (n=3) for each genotype. A Mann-Whitney test was used to analyze significance (****, *p*-value <0.0001). Scale bar = 10 μm

### Nab2 is required in neurons to restrict lipid droplet size in the fat body

Nab2 plays an important role in neuronal function to support viability, learning/memory and to guide axon projection (Pak et al. 2011, Lancaster et al. 2024). A subset of neurons found in the brain are the IPCs which are the only cells that produce the Dilp2 and Dilp5 peptides (Géminard et al. 2009). Given that increased *dilp2* and *dilp5* mRNA in the *Nab2^null^* brains correlates with increased lipid droplet size in the *Nab2^null^* fat body, we investigated whether Nab2 is required within specific cells to regulate fat body lipid droplet size. These experiments take advantage of the *Drosophila* model using the Gal4/UAS system (Brand et al. 1993) to drive a *Nab2* RNAi transgene (Dietzl et al. 2007) that depletes *Nab2* specifically in specific cell types, including neurons, IPCs and glia. Glia supply neighboring neurons with nutrients and modulate synaptic signaling (Uchewa et al. 2025). Therefore, a possible role for Nab2 in glia cells was investigated in parallel with neurons and IPCs. We first tested whether Nab2 is required in neurons to regulate lipid droplet size in the fat body (Pak et al. 2011). As shown in Figure 4A, Nab2 depletion in all neurons (*C155Gal4* driver) yields fat bodies with an average lipid droplet size that is similar to *Nab2^null^* female larvae demonstrating that depletion of Nab2 in all neurons is sufficient to elevate lipid storage in the fat body. Next, we tested whether RNAi-mediated depletion of Nab2 in either the IPCs (*Dilp2Gal4* driver), which are the neuronal cells that are the source of Dilp2/5 production (Géminard et al. 2009), or glia (*RepoGal4* driver) is sufficient to drive lipid accumulation in the fat body. Neither depletion of Nab2 from the IPCs (Figure 4B) nor glia cells (Figure 4C) caused a statistically significant change in lipid droplet size in the fat body. These experiments provide evidence that Nab2 is required in neurons to regulate lipid droplet size in the fat body.

To test whether restoring Nab2 expression only in neurons is sufficient to restore proper lipid droplet size in *Nab2^null^* female larvae, we expressed a *UAS-Nab2-FLAG* transgene in the neurons (*C155Gal4* driver) of *Nab2^null^* female larvae creating larvae with Nab2 absent in all cells except neurons. Expressing *Nab2-FLAG* in the neurons of these *Nab2^null^* female larvae produces a statistically significant rescue of the increased fat body lipid droplet phenotype (Figure 4A). In sum, these Gal4/UAS data indicate that Nab2 is both necessary and sufficient in neurons to regulate lipid droplet size in the female larval fat body.

### Depletion of dilp2/dilp5 rescues increased lipid droplet size in Nab2^null^ female larval fat body

The increase in the steady-state level of *dilp2* and *dilp5* transcripts in *Nab2^null^* females supports a model in which Nab2 loss elevates Dilp production in the IPCs, which in turn drives excess lipid storage in the female fat body. However, elevated *dilp2* and *dilp5* mRNAs in the *Nab2^null^*females may not be causally linked to the increased size of fat body lipid droplets. To assess whether the increase in *dilps* is required for increased fat body lipid droplet size, we used RNAi to deplete either *dilp2* or *dilp5* in the IPCs (*Dilp2Gal4* driver) in *Nab2^null^* females and assessed lipid droplet size in the fat body. As shown in Figure 5A and 5B, IPC-specific depletion of either *dilp2* or *dilp5* is sufficient to restore the *Nab2^null^*female lipid droplet size to sizes similar to those observed in the fat bodies of Control females. This finding confirms that *dilp2* and *dilp5* are required for the female-specific lipid droplet defect in *Nab2^null^*female larvae and provides evidence that Nab2 acts upstream of the IPCs to regulate *dilp2* and *dilp5* mRNA levels and lipid storage in females.

**FIGURE 5.**
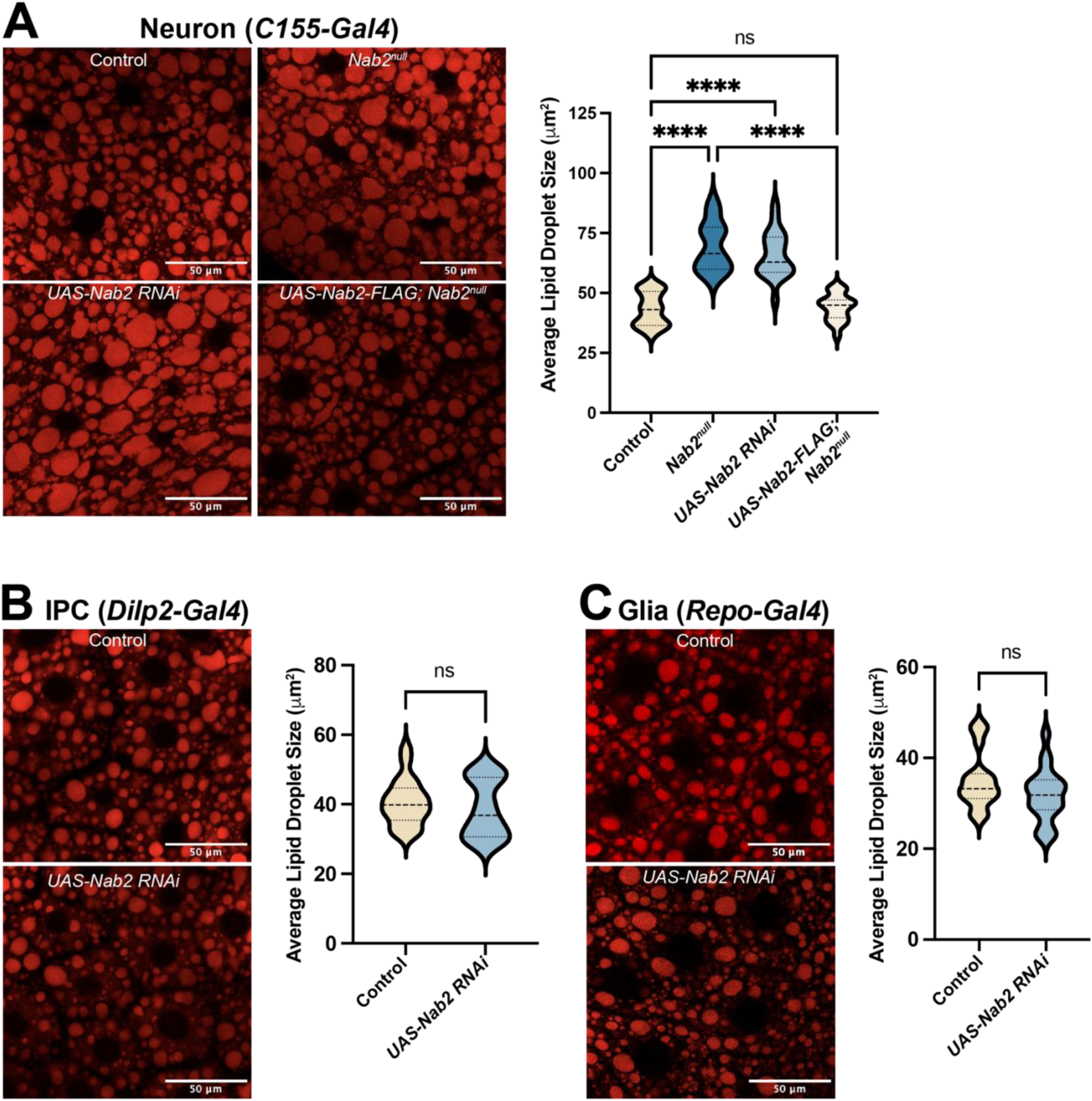
Nab2 is required in neurons to restrict lipid droplet size in the fat body. (A) Lipid droplet size measurements using *C155Gal4* driver (neurons). Representative fields are shown for Control (*C155Gal4*), *Nab2^null^* (*C155Gal4; Nab2^null^*), RNAi-mediated depletion of *Nab2* (*C155Gal4; UAS-Nab2-RNAi*) and neuronal rescue via UAS-*Nab2*-FLAG in *Nab2^null^* background (*C155Gal4; UAS-Nab2-FLAG; Nab2^null^*). (B) Lipid droplet size measurements using *Dilp2Gal4* driver (IPCs) to express UAS-*Nab2* RNAi in IPCs. Representative fields are shown for Control (*Dilp2Gal4*) and RNAi-mediated depletion of *Nab2* (*Dilp2Gal4; UAS-Nab2-RNAi*). (C) Lipid Droplet size using *RepoGal4* driver (Glia) to express UAS-*Nab2* RNAi in Glia. Representative fields are shown for Control (*RepoGal4*) and RNAi-mediated depletion of *Nab2* (*RepoGal4; UAS-Nab2-RNAi*). For quantitation, 20 age matched biological replicates were used for each genotype. One-way ANOVA or Welches t-test was performed and significance values are indicated. (****, *p*-value <0.0001). Scale bar = 50 μm

## Discussion

Here, we identify a previously uncharacterized role for the RBP *Drosophila* Nab2 as a female-specific regulator of fat storage in flies. GO analysis revealed various metabolic biological processes and lipid biological processes that are significantly enriched in differential gene analysis from *Nab2^null^* female and male adult heads compared to sex-matched Controls. Loss of Nab2 leads to an increase in the levels of *dilp2* and *dilp5* transcripts in female larvae. As Dilp2 and Dilp5 are produced and released by the IPCs in the brain to promote fat storage in peripheral tissues such as the *Drosophila* fat body (Garofalo 2002, Géminard et al. 2009), these results suggest that Nab2 function is required in neurons to regulate *dilp* production and insulin signaling as loss of Nab2 leads to elevate lipid storage in the fat body. Given that *Nab2^null^* female larvae show increased size but no change in food consumption, evidence supports a molecular role for Nab2 in regulating insulin signaling and fat storage rather than a role of Nab2 in controlling feeding behavior that results in increased levels of *dilp2* and *dilp5* transcript. Consistent with this model, Nile Red staining reveals a significant increase in the size of the lipid droplets found in the *Nab2^null^* female larval fat body compared to Control, but no such phenotype in males. Further investigation of the lipid droplet phenotype led to the conclusion that loss of Nab2 specifically in neurons leads to increased lipid droplet size in the female fat body. However, loss of Nab2 in IPCs and glia cells caused no change in lipid droplet size. Importantly, expressing *Nab2* solely in the neurons of a *Nab2^null^* female rescues the lipid droplet phenotype seen in *Nab2^null^* female fat body. Together, these observations support a model (Figure 6) for Nab2 in which its role in a neuronal population within this neurometabolic circuit regulates *dilp* production in the IPCs to regulate lipid storage in the *Drosophila* fat body.

**FIGURE 6.**
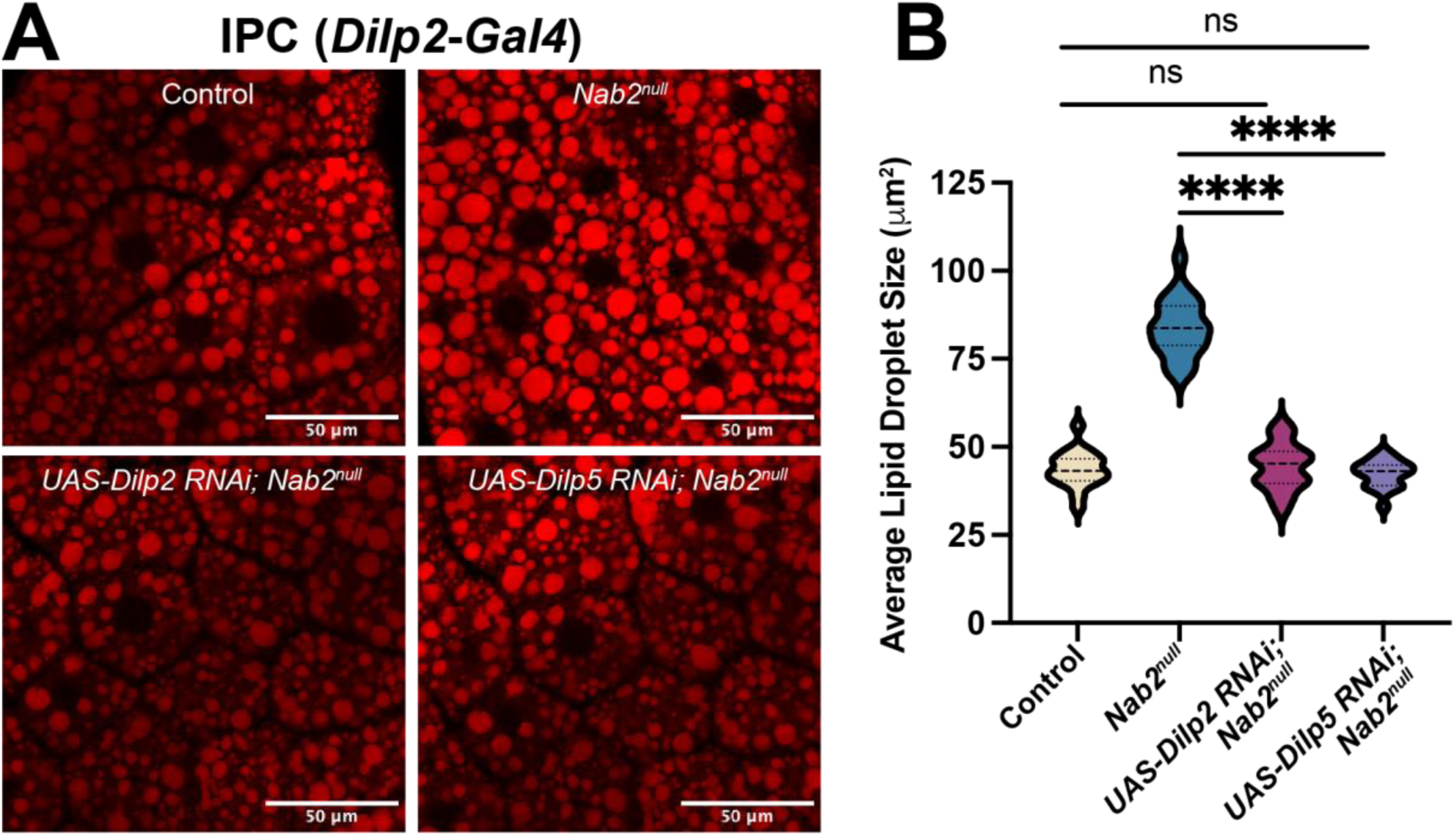
Depletion of dilp2/dilp5 rescues increased lipid droplet size in Nab2^null^ female larval fat body. (A) Representative 3^rd^ instar female larval fat body of the indicated genotypes stained by Nile Red. (B) Lipid droplet size measurements using *Dilp2Gal4* driver (IPCs). Representative fields are shown for Control (*Dilp2Gal4*), *Nab2^null^* (*Dilp2Gal4; Nab2^null^*), RNAi mediated depletion of *Dilp2* (*Dilp2Gal4/UAS-Dilp2-RNAi; Nab2^null^*) or *Dilp5* (*Dilp2Gal4/UAS-Dilp5-RNAi; Nab2^null^*) in the background of *Nab2^null^* larvae. For each genotype, 20 age-matched biological replicates were used. One-way ANOVA was performed and significance values are indicated. (****, *p*-value <0.0001). Scale bar = 50 μm

**FIGURE 7.**
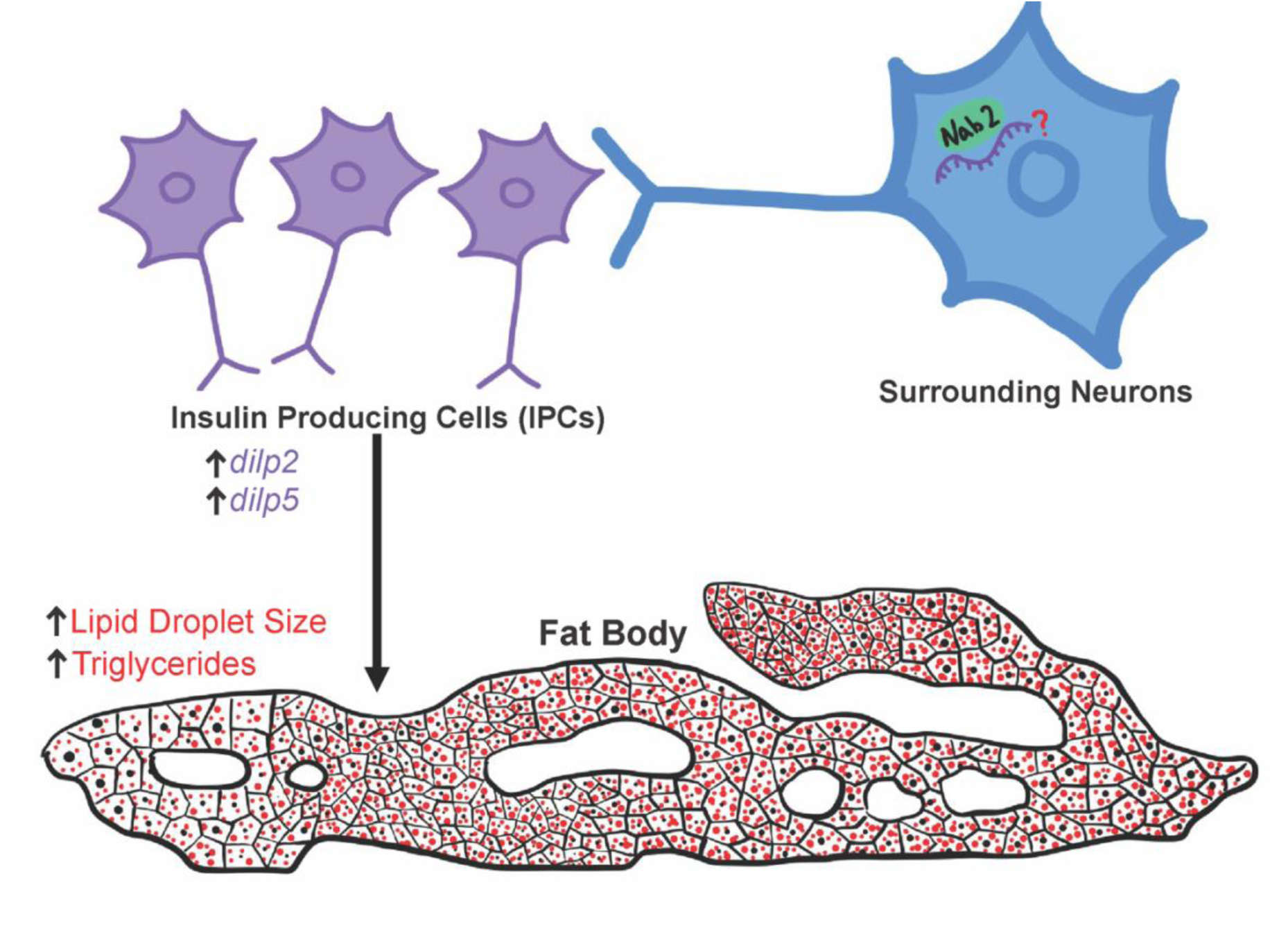
Proposed model for Nab2 regulation of *dilp2/dilp5* transcript levels and lipid storage in female larvae. Loss of Nab2 in neurons, but not the insulin producing cells (IPCs), increases the steady state levels of *dilp2* and *dilp5* transcripts in the brain, resulting in increased fat body lipid droplet size and TAG levels in females, suggesting that Nab2 regulates systemic lipid homeostasis by modulation of insulin-like peptide levels through a subset of neurons surrounding the IPCs. The model proposes that Nab2 regulates a target RNA in a specific subset of neurons that has a role in the insulin signaling pathway in the brain involved in Dilp production/secretion to signal lipid storage in the fat body.

Sex-specific differences in both *Drosophila* and humans involve male and female differences in hormone signaling, nutrient utilization and fat storage (Varlamov et al. 2014, Dähn et al. 2025). Specifically, females exhibit distinct patterns of lipid accumulation and nutrient storage associated with development and reproductive demands that can influence insulin signaling and energy homeostasis (Varlamov et al. 2014, Kim et al. 2021). Our data suggest that Nab2 maybe a key part of this mechanism in flies via a role in the female brain. Loss of Nab2 may influence how energy usage and storage occurs in females possibly due to differences in developmental insulin signaling compared to males. These data support a role for Nab2 in female-specific metabolic regulation involving insulin signaling and lipid storage.

Previous studies have established a role for Nab2 in regulating neurons in both larval and adult *Drosophila* (Corgiat et al. 2022, Lancaster et al. 2024). Interestingly, neuronal depletion of Nab2 recapitulated the lipid droplet phenotype seen in the *Nab2^null^* female fat body but IPC depletion of Nab2 did not. These data support a role for Nab2 in the neuronal cells that are upstream of the IPCs to regulate Dilp production as shown in Figure 6. Importantly, several neuronal circuits communicate with the IPCs to control Dilp production and secretion in response to various developmental and environmental cues (Kapan et al. 2012, Oh et al. 2019, Bisen et al. 2025). Consistent with the idea that increased lipid droplet size in the fat body is linked to elevated *dilp2* and *dilp5* transcript levels in *Nab2^null^* female larval brain, RNAi-mediated depletion of either *dilp2* or *dilp5* in the IPCs of a *Nab2^null^* female larva restores lipid droplet size to Control levels. While Dilp2 and Dilp5 have distinct developmental roles (Henstridge et al. 2026), these results are consistent with elevated *dilp2* and *dilp5* driving excess lipid storage in *Nab2^null^*female larvae. The results support a model (Figure 6) where Nab2 functions within a subset of neurons that act upstream of the IPCs and influence the expression and possibly secretion of Dilp2 and Dilp5. Therefore, loss of Nab2 in this yet unidentified set of neurons leads to increased levels of both *dilp2* and *dilp5* transcript, which results in increased lipid storage in the fat body. Further studies focused on identifying the direct mRNA target of Nab2 in these candidate neuronal subsets will be important for understanding the post-transcriptional role for Nab2 in regulating this neurometabolic pathway. One possible target of Nab2 that could be involved in female-specific metabolism is the sex determination gene Sex lethal (*sxl*) (Penalva et al. 2003). However, Sxl has only been shown to control lipid metabolism from within fat body cells (Diaz et al. 2023). Notably, a recent study identified the human protein Death-inducer obliterator 1 (DIDO1) as a sex-specific determinant of obesity in females (Kaisinger et al. 2023). The *Drosophila* orthologue of DIDO1 is protein partner of snf (Pps) which is part of the machinery that is required for the sex-specific splicing of *Sxl* (Johnson et al. 2010), further implicating the *sxl* transcript as a possible Nab2 target in female metabolic regulation. Past research shows that Nab2 regulates the splicing of *sxl* mRNA in brain neurons and in the absence of Nab2 there is a masculinization of Nab2 mutant females (Jalloh and Lancaster et al. 2023). Importantly, restoring *sxl* splicing can rescue many of the phenotypes exhibited in the Nab2^null^ flies (Jalloh and Lancaster et al. 2023).

Our findings contribute to the growing evidence that loss of a single RBP can cause both neurological disorders and metabolic dysfunction. This work suggests that metabolic dysfunction in individuals with a neurological disorder may be a consequence of post-transcriptional gene regulation caused by loss of a single RBP. Uncovering these connections between RBP loss and metabolic dysfunction is important as there can be elevated health concerns for individuals with neurological disorders (Feigin et al. 2020). Also, the recent discovery of genetic variants that cause sex-specific obesity (Kaisinger et al. 2023) supports the need for further research to understand how loss of RBPs that have previously been linked to neurological disease may lead to sex-specific defects in metabolic regulation. Specifically, understanding the role of Nab2/ZC3H14 in metabolic homeostasis may provide new insight into the metabolic complications found in patients with neurological disorders possibly revealing a conserved mechanism by which RBPs with important roles in the nervous system regulate insulin signaling and lipid metabolism.

## Materials and Methods

### Resources Table

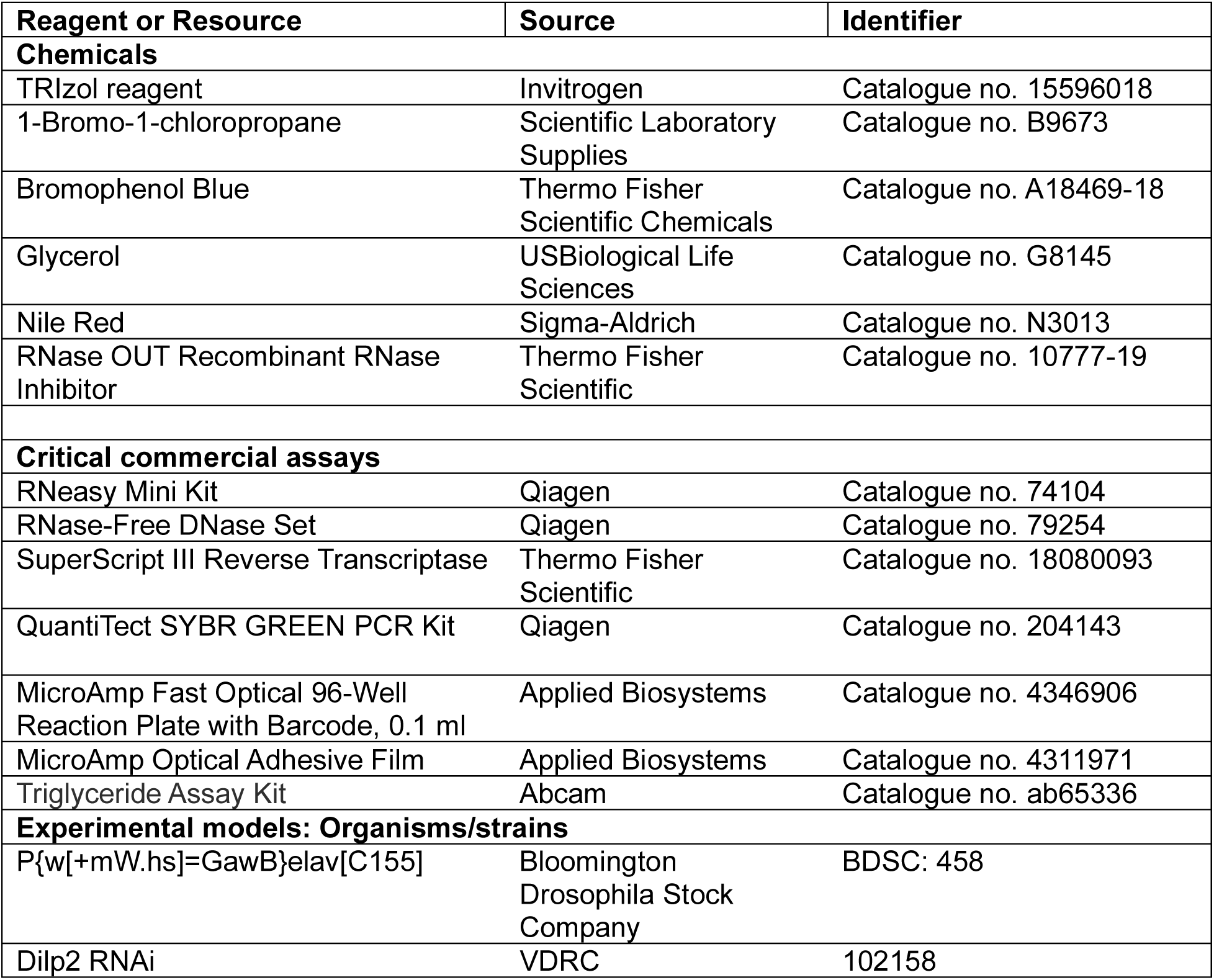
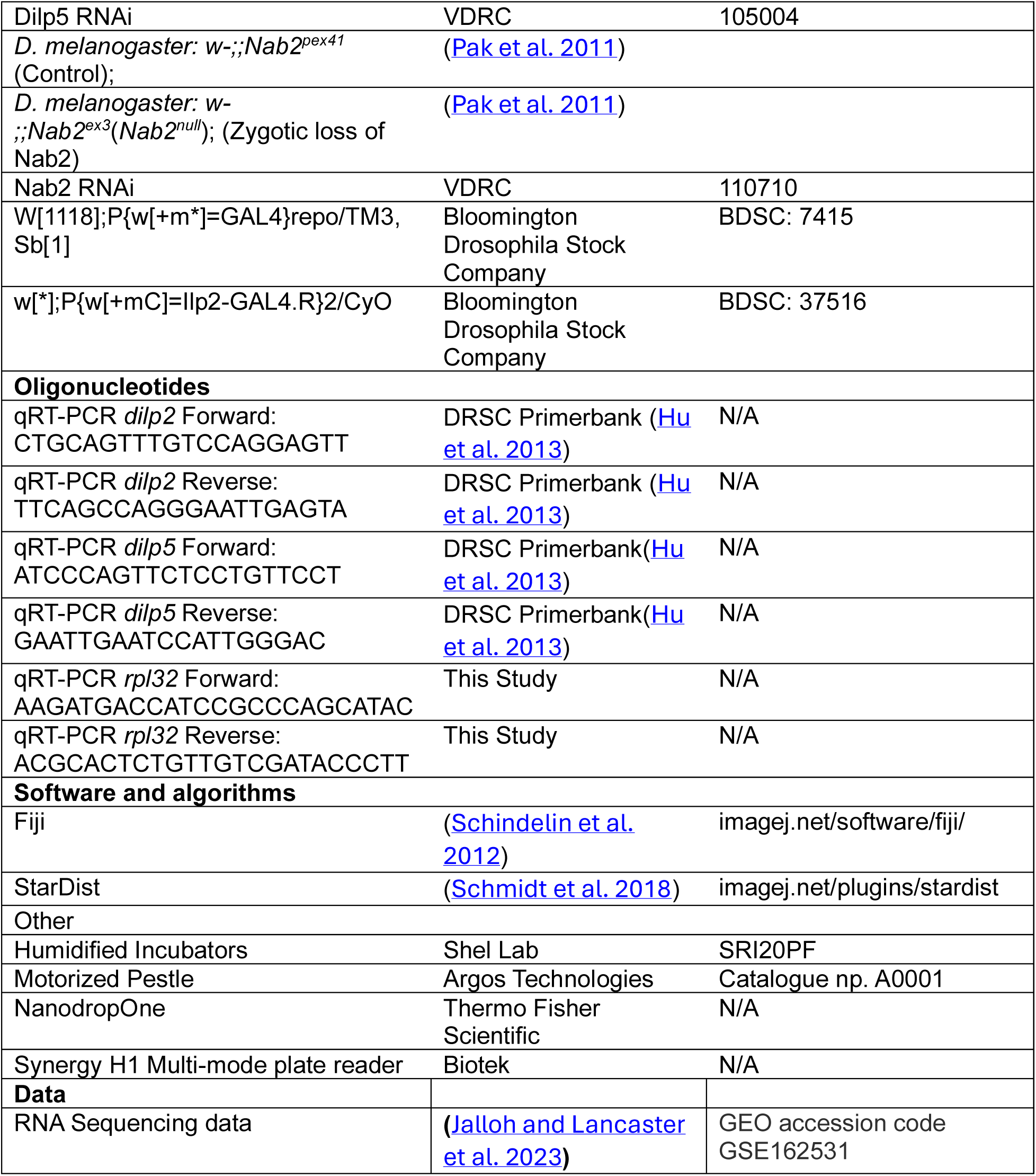

## Resource availability

### Lead Contact

Further information and requests for resources and reagents should be directed to and will be fulfilled by the Lead Contact, Ken Moberg.

### Materials availability

The *D. melanogaster* lines generated in this study are available by contacting the Lead Contact.

## Method Details

### D. melanogaster stocks and genetics

*D. melanogaster* stocks were raised on standard cornmeal agar and maintained in humidified incubators at 25°C with 12-h light/dark cycles. Crosses were reared under the same conditions and supplemented with dry yeast. The GAL4-UAS binary transgenic system was used to express transgenes of interest. Details of genotypes used in this study are described in the Key Resources Table.

### RNA Sequencing Analysis

Raw read FASTA files were analyzed from (Jalloh and Lancaster et al. 2023) GEO accession code GSE162531. Adapter sequences were trimmed using Trim Galore (Krueger, F. (2026). Trim Galore (Version 2.2.0) [Computer software]. https://github.com/FelixKrueger/TrimGalore), followed by alignment to the *Drosophila melanogaster* reference transcriptome (NCBI assembly database accession number GCA_0000012154) using STAR (v. 2.7.11b). RSEM (v1.3.3) was used to quantify gene-level mRNA expression. Differential gene expression analysis was performed using the DESeq2 (v1.50.2) package in R. Genes with an adjusted *p*-value < 0.05 were considered significantly differentially expressed.

### Gene ontology analysis

*Gene* Ontology (GO) analysis was performed using the clusterProfiler package in R to characterize the differentially expression genes identified. Reference GO gene sets (org.Dm.eg.db) were used to determine the top biological processes and lipid processes GO terms upon loss of Nab2 in females and males compared to sex-matched Controls. Statistical significance was assessed and GO terms with an adjusted *p*-value <0.05 were considered significantly enriched.

### RNA isolation for real-time qPCR

Total RNA was isolated from ∼20 adult fly heads or ∼25 larval brains using the TRIzol (Invitrogen) method. *Drosophila* adult heads and larval brains were homogenized in 0.1ml TRIzol using a motorized pestle (Argos Technologies) on ice. TRIzol was added to sample to bring to a total volume of 0.5 ml and 0.1 ml of 1-Bromo-3-chloropropane (Scientific Laboratory Supplies) was added. Samples were vortexed on high speed for 20 s and incubated at room temperature for 3 min. Next, samples were centrifuged for 15 min at 12,000 x g at 4 °C. The top aqueous phase was transferred into a clean Eppendorf tube. An equal volume of 100% RNA-free ethanol was added to each sample. Samples were inverted 10 times. Samples were loaded onto a RNeasy column seated in a collection tube from the RNeasy mini kit (Qiagen). Then samples were centrifuged for 30 sec at 8,000 x g and the flow-through was discarded. Next, 350 μl Buffer RW1 was loaded onto the RNeasy spin column, centrifuged for 15 s at 8000 x g to wash the spin column membrane and flow-through was discarded. DNase Master Mix (10 ul DNase I and 70 ul Buffer RDD) was then added directly to the RNeasy spin column membrane and incubate at RT for 15 min followed by the addition of 350 μl Buffer RW1 to the RNeasy spin column. Samples were centrifuged for 15 s at 8000 x g and flow-through was discarded. Samples columns were then transferred into a new collection tube. Then, 500µl buffer RPE was added to the column, centrifuged for 30 sec at 8,000 x g and the flow-through was discarded. 500µl buffer RPE was added to the sample, sample was centrifuged for 2 min at 8,000 x g and flow-through was discarded. The column was spun for 1 min at 8,000 x g to get rid of remaining buffer in the column. The column was transferred to a new 1.5 mL collection tube and 15 µL of RNase-free water directly onto the column membrane. Samples were incubated at room temperature for 2 min and then spin 1 min at 8,000 x g to elute RNA. The elution step was repeated with the same 1.5mL collection tube, generating 30 µL of RNA in total. Primers used for all PCR reactions are listed in the Key Resources Table.

### *Drosophila* decapitation

Flies were collected and frozen at -80°C. Frozen flies were placed on a metal tray over dry ice. Forceps were placed between the head and thorax to remove the head. Heads were placed in Eppendorf tubes, on dry ice before proceeding with RNA extraction.

### *Drosophila* larval brain dissection

For *Drosophila* larval brain RNA extraction, third instar larval brains were dissected in PBS. Larvae were transferred to a drop of PBS on a dissecting mat. A pair of forceps was used to grasp the mid region of a single larva. Dissecting scissors were used to cut the larvae right above the forceps at the mid region. Forceps were used to grasp the anterior mouth hooks. Another set of forceps was used to roll the cuticle up the first pair of forceps that are grasping the mouth hooks so that the cuticle is turned inside out. With the brain now exposed other tissues were disregarded from the rest of the larvae and the brain was transferred to an Eppendorf tube containing 0.1 ml of TRIzol.

### Larval weight

Third instar larvae were collected and briefly washed in ddH2O to remove any food particles that remained on the outside of the larvae. Larvae were gently blotted with a dry KimWipe and then 10 larvae were placed into a clean Eppendorf tube, and group weight was measured using a precision microbalance.

### Larval Size

Third instar larvae were collected and briefly washed in ddH2O to remove any food particles that remained on the outside of the larvae. Larvae were gently blotted with a dry KimWipe. Larvae were then place on a frozen metal block to decrease larval movement during imaging. A ruler was placed at the top of each image to allow for accurate larval size measurements using Fiji/ImageJ. Measurements were taken from the most anterior end of the larvae to the most posterior end to gather larval length measurements. To measure the width of the larvae the midline each larva was measured.

### Larval Food Steady State Levels

*Drosophila* adults were allowed to lay on petri dishes filled with food supplemented with bromophenol blue (often used for consumption assays in *Drosophila* (Walls et al. 2013)) for 2 hrs. Larvae were raised until early third instar larvae as feeding decreases sharply in later third instar phases. Early third instar larvae were collected and briefly washed in ddH2O to remove any food particles that reminded on the outside of the larvae. Larvae were gently blotted with a dry KimWipe. 4 larvae were placed into a clean Eppendorf tube. 0.2 ml of ddH2O was added to the Eppendorf tube and larvae were homogenized using a motorized pestle (Argos Technologies) for 2 min. Tubes were briefly centrifuged for 15 sec to pellet any debris. 50 μl of supernatant was added to 150 μl of ddH2O in a single well on a 96 well plate. This was repeated two more times for each sample to provide triplicate measurements for each sample on the plate. For the blank on the plate, 50 μl of homogenate from ∼4 non-blue larvae was mixed with 150 μl of ddH2O in a well. To quantify food consumption standards of bromophenol blue were used. Through serial dilution the following standards were created to quantify the amount of food extracted from larvae: 125 μg/ml, 62.5 μg/ml. 31.25 μg/ml, 15.62 μg/ml and 7.8125 μg/ml. The absorbance was measured using a Synergy H1 multi-mode plate reader to measure the absorbance at 595nm. Using the standard curve the concentration of dye in each diluted sample was calculated which was then converted to volume of food consumed. This value was then divided by 4 to get an approximate measurement for food per larvae in a sample.

### Nile Red Staining

The fat body from third instar larvae was dissected in 1xPBS and then fixed in 4% PFA at room temperature for 30min. Samples were washed twice for 5 min in 1xPBS. Then samples were incubated with Nile Red (stock 1000μg/ml, diluted 1:20 in 1xPBS) for 20 min. Samples were washed with ddH2O twice for 5 min. Samples were mounted using 75% glycerol.

### Nile Red Image Analysis

To quantify lipid droplet size in third instar larval fat body, Nile Red confocal images captured on a Nikon A1R inverted Confocal microscope were analyzed using the StarDist plugin in Fiji/ImageJ. StarDist enables accurate segmentation of round or elliptical objects.

### Triacylglyceride Measurements

To quantify triacylglyceride (TAG) levels in the 3^rd^ instar *Drosophila* larvae a TAG plate reader kit (Abcam Catalogue no. ab65336) was used. Upon collection 3^rd^ instar age matched larvae were cleaned with 1 X PBS and dried on a KimWipe to remove excess food. Four age matched 3^rd^ instar larvae for Control and *Nab2^null^* were obtained for each biological replicate (n=5) for both males and females in an Eppendorf tube. Larvae were homogenized in 400μl of 5% NP-40 and tissue sample prep was followed as outlined in the kit protocol. 5μl of supernatant was used to determine the level of TAG in the sample for the colorimetric assay in the kit using Synergy H1 multi-mode plate reader at OD 570 nm. 1μl of supernatant was used to measure the amount of protein in the sample using a BSA assay. Samples were read at an OD 595 nm using Beckman DU 530 UV/Vis spectrophotometer.

### Hemolymph collection and lipid measurement

Ten (10) late wandering stage larva were collected and rinsed 3x in water to thoroughly remove residual food, then placed on filter paper to remove exterior moisture. The larvae were then transferred to a well of a dry glass collection dish pre-chilled on ice. Forceps were used to tear the larval cuticle open midway down the body and allowed to bleed out. Yield is <1μl per animal. A 5μl pipette fitted with 2μl filter tips was used to remove the pooled hemolymph around the corpses without damaging exposed tissues and organs. Nile Red (1000μg/ml) was added at a 1:20 dilution and the samples were spotted onto a slide, mounted with a coverslip and imaged on a Leica CCD camera with Cy3/Rhodamine excitation. The yield is <1μl per animal. For analysis, images were imported into Fiji and individual droplets were defined by segmenting based on >0.5 circularity. Data were imported into Prism for analysis. A Mann-Whitney test was used to analyze significance of the number/size distribution between three biological replicates of data.

### Statistical analysis

Group analysis on biological triplicate experiments was performed using welches t-test, Mann-whitney test or one-way Anova (turkeys multiple comparison test) on GraphPad (Prism). Sample sizes (*n*) and *p*-values are denoted in text, figures and /or figure legends. *p*-values are indicated by asterisks (e.g., *, *p* < 0.05)

## Abreviations

ZC3H14: zinc finger Cys-Cys-Cys-His-type containing 14
Nab2: nuclear polyadenosine RNA-binding protein 2
RBP: RNA-binding protein
Dilp: *Drosophila* insulin-like peptide
TAG: Triacylglycerides
IPCs: Insulin Producing Cells

## Acknowledgements

Research reported in this publication was supported by NINDS/NIH awards R01-NS125768 to AHC and KHM, and F31-NS135980 to JNG. DUG was supported by NICHD/NIH award R01-HD102534. RMM was supported by a T32 graduate training award from NIGMS (GM149422). The content is solely the responsibility of the authors and does not necessarily represent the official views of the National Institutes of Health.

## Notes

### Competing Interest Statement

The authors have declared no competing interest.

## References

Ahima, R. S. and D. A. Antwi (2008). “Brain regulation of appetite and satiety.” Endocrinol Metab Clin North Am 37(4): 811–823.

Aibara, S., J. M. Gordon, A. S. Riesterer, S. H. McLaughlin and M. Stewart (2017). “Structural basis for the dimerization of Nab2 generated by RNA binding provides insight into its contribution to both poly(A) tail length determination and transcript compaction in Saccharomyces cerevisiae.” Nucleic Acids Res 45(3): 1529–1538.

Alpert, T., K. Straube, F. Carrillo Oesterreich, L. Herzel and K. M. Neugebauer (2020). “Widespread Transcriptional Readthrough Caused by Nab2 Depletion Leads to Chimeric Transcripts with Retained Introns.” Cell Rep 33(4): 108324.

Arrese, E. L. and J. L. Soulages (2010). “Insect fat body: energy, metabolism, and regulation.” Annu Rev Entomol 55: 207–225.

Awari, A., D. Kaushik, A. Kumar, E. Oz, K. Çadırcı, C. Brennan, C. Proestos, M. Kumar and F. Oz (2025). “Obesity Biomarkers: Exploring Factors, Ramification, Machine Learning, and AI-Unveiling Insights in Health Research.” MedComm (2020) 6(7): e70169.

Banerjee, D. and A. Mani (2025). “Obesity’s systemic impact: exploring molecular and physiological links to diabetes, cardiovascular disease, and heart failure.” Front Endocrinol (Lausanne) 16: 1681766.

Bardoni, B., S. Abekhoukh, S. Zongaro and M. Melko (2012). “Intellectual disabilities, neuronal posttranscriptional RNA metabolism, and RNA-binding proteins: three actors for a complex scenario.” Prog Brain Res 197: 29–51.

Benegiamo, G., L. S. Mure, G. Erikson, H. D. Le, E. Moriggi, S. A. Brown and S. Panda (2018). “The RNA-Binding Protein NONO Coordinates Hepatic Adaptation to Feeding.” Cell Metab 27(2): 404–418.e407.

Beshel, J., J. Dubnau and Y. Zhong (2017). “A Leptin Analog Locally Produced in the Brain Acts via a Conserved Neural Circuit to Modulate Obesity-Linked Behaviors in Drosophila.” Cell Metab 25(1): 208–217.

Bisen, R. S., F. M. Iqbal, F. Cascino-Milani, T. Bockemühl and J. M. Ache (2025). “Nutritional state-dependent modulation of insulin-producing cells in Drosophila.” Elife 13.

Biswas, P., J. A. Bako, J. B. Liston, H. Yu, L. W. Wat, C. J. Miller, M. D. Gordon, T. Huan, M. Stanley and E. J. Rideout (2025). “Insulin/insulin-like growth factor signaling pathway promotes higher fat storage in Drosophila females.” bioRxiv.

Brand, A. H. and N. Perrimon (1993). “Targeted gene expression as a means of altering cell fates and generating dominant phenotypes.” Development 118(2): 401–415.

Brogiolo, W., H. Stocker, T. Ikeya, F. Rintelen, R. Fernandez and E. Hafen (2001). “An evolutionarily conserved function of the Drosophila insulin receptor and insulin-like peptides in growth control.” Curr Biol 11(4): 213–221.

Broughton, S., N. Alic, C. Slack, T. Bass, T. Ikeya, G. Vinti, A. M. Tommasi, Y. Driege, E. Hafen and L. Partridge (2008). “Reduction of DILP2 in Drosophila triages a metabolic phenotype from lifespan revealing redundancy and compensation among DILPs.” PLoS One 3(11): e3721.

Ceron-Codorniu, M., P. Torres, A. Fernàndez-Bernal, S. Rico-Rios, J. C. Serrano, M. P. Miralles, M. Beltran, A. Garcera, R. M. Soler, R. Pamplona and M. Portero-Otín (2024). “TDP-43 dysfunction leads to bioenergetic failure and lipid metabolic rewiring in human cells.” Redox Biol 75: 103301.

Cesur, M. F., A. Basile, K. R. Patil and T. Çakır (2023). “A new metabolic model of Drosophila melanogaster and the integrative analysis of Parkinson’s disease.” Life Sci Alliance 6(8).

Chatterjee, N. and N. Perrimon (2021). “What fuels the fly: Energy metabolism in Drosophila and its application to the study of obesity and diabetes.” Sci Adv 7(24).

Church, R. B. and F. W. Robertson (1966). “A biochemical study of the growth of Drosophila melanogaster.” Journal of Experimental Zoology 162(3): 337–351.

Conlon, E. G. and J. L. Manley (2017). “RNA-binding proteins in neurodegeneration: mechanisms in aggregate.” Genes Dev 31(15): 1509–1528.

Cooper, T. A., L. Wan and G. Dreyfuss (2009). “RNA and disease.” Cell 136(4): 777–793.

Corbett, A. H. (2018). “Post-transcriptional regulation of gene expression and human disease.” Curr Opin Cell Biol 52: 96–104.

Corgiat, E. B., S. M. List, J. C. Rounds, A. H. Corbett and K. H. Moberg (2021). “The RNA-binding protein Nab2 regulates the proteome of the developing Drosophila brain.” J Biol Chem 297(1): 100877.

Corgiat, E. B., S. M. List, J. C. Rounds, D. Yu, P. Chen, A. H. Corbett and K. H. Moberg (2022). “The Nab2 RNA-binding protein patterns dendritic and axonal projections through a planar cell polarity-sensitive mechanism.” G3 (Bethesda) 12(6).

Dähn, S. and A. E. Wagner (2025). “Drosophila melanogaster as a model organism to investigate sex specific differences.” Sci Rep 15(1): 19648.

Darnell, R. B. (2013). “RNA protein interaction in neurons.” Annu Rev Neurosci 36: 243–270.

DiAngelo, J. R. and M. J. Birnbaum (2009). “Regulation of fat cell mass by insulin in Drosophila melanogaster.” Mol Cell Biol 29(24): 6341–6352.

Diaz, A. V., D. Stephenson, T. Nemkov, A. D’Alessandro and T. Reis (2023). “Spenito-dependent metabolic sexual dimorphism intrinsic to fat storage cells.” Genetics 225(3).

Dietzl, G., D. Chen, F. Schnorrer, K. C. Su, Y. Barinova, M. Fellner, B. Gasser, K. Kinsey, S. Oppel, S. Scheiblauer, A. Couto, V. Marra, K. Keleman and B. J. Dickson (2007). “A genome-wide transgenic RNAi library for conditional gene inactivation in Drosophila.” Nature 448(7150): 151–156.

Efeyan, A., W. C. Comb and D. M. Sabatini (2015). “Nutrient-sensing mechanisms and pathways.” Nature 517(7534): 302–310.

Fasken, M. B., A. H. Corbett and M. Stewart (2019). “Structure-function relationships in the Nab2 polyadenosine-RNA binding Zn finger protein family.” Protein Sci 28(3): 513–523.

Feigin, V. L., T. Vos, E. Nichols, M. O. Owolabi, W. M. Carroll, M. Dichgans, G. Deuschl, P. Parmar, M. Brainin and C. Murray (2020). “The global burden of neurological disorders: translating evidence into policy.” Lancet Neurol 19(3): 255–265.

Fujimoto, T. and R. G. Parton (2011). “Not just fat: the structure and function of the lipid droplet.” Cold Spring Harb Perspect Biol 3(3).

Gabs, E., E. Aalto-Setälä, A. Välisaari, A. M. Malinen, T. H. Jensen, S. H. McLaughlin, L. A. Passmore and M. Turtola (2025). “A kinetic ruler controls mRNA poly(A) tail length.” Genes Dev 39(21-22): 1377–1394.

Garofalo, R. S. (2002). “Genetic analysis of insulin signaling in Drosophila.” Trends Endocrinol Metab 13(4): 156–162.

Géminard, C., E. J. Rulifson and P. Léopold (2009). “Remote control of insulin secretion by fat cells in Drosophila.” Cell Metab 10(3): 199–207.

Glisovic, T., J. L. Bachorik, J. Yong and G. Dreyfuss (2008). “RNA-binding proteins and post-transcriptional gene regulation.” FEBS Lett 582(14): 1977–1986.

Graham, P. and L. Pick (2017). “Drosophila as a Model for Diabetes and Diseases of Insulin Resistance.” Curr Top Dev Biol 121: 397–419.

Greenspan, P., E. P. Mayer and S. D. Fowler (1985). “Nile red: a selective fluorescent stain for intracellular lipid droplets.” J Cell Biol 100(3): 965–973.

Grenier St-Sauveur, V., S. Soucek, A. H. Corbett and F. Bachand (2013). “Poly(A) tail-mediated gene regulation by opposing roles of Nab2 and Pab2 nuclear poly(A)-binding proteins in pre-mRNA decay.” Mol Cell Biol 33(23): 4718–4731.

Guthrie, C. R., L. Greenup, J. B. Leverenz and B. C. Kraemer (2011). “MSUT2 is a determinant of susceptibility to tau neurotoxicity.” Hum Mol Genet 20(10): 1989–1999.

Guthrie, C. R., G. D. Schellenberg and B. C. Kraemer (2009). “SUT-2 potentiates tau-induced neurotoxicity in Caenorhabditis elegans.” Hum Mol Genet 18(10): 1825–1838.

Hagerman, R. J., E. Berry-Kravis, H. C. Hazlett, D. B. Bailey, Jr., H. Moine, R. F. Kooy, F. Tassone, I. Gantois, N. Sonenberg, J. L. Mandel and P. J. Hagerman (2017). “Fragile X syndrome.” Nat Rev Dis Primers 3: 17065.

He, S., E. Valkov, S. Cheloufi and J. Murn (2023). “The nexus between RNA-binding proteins and their effectors.” Nat Rev Genet 24(5): 276–294.

Henstridge, M. A., B. Slater, J. R. Kannangara and C. K. Mirth (2026). “Insulin-like peptides play distinct roles in nutrient-dependent plasticity in Drosophila.” G3 Genes|Genomes|Genetics 16(2).

Hu, Y., R. Sopko, M. Foos, C. Kelley, I. Flockhart, N. Ammeux, X. Wang, L. Perkins, N. Perrimon and S. E. Mohr (2013). “FlyPrimerBank: an online database for Drosophila melanogaster gene expression analysis and knockdown evaluation of RNAi reagents.” G3 (Bethesda) 3(9): 1607–1616.

Jalloh, B., C. L. Lancaster, J. C. Rounds, B. E. Brown, S. W. Leung, A. Banerjee, D. J. Morton, R. S. Bienkowski, M. B. Fasken, I. J. Kremsky, M. Tegowski, K. Meyer, A. Corbett and K. Moberg (2023). “The Drosophila Nab2 RNA binding protein inhibits m(6)A methylation and male-specific splicing of Sex lethal transcript in female neuronal tissue.” Elife 12.

Jo, M., S. Lee, Y. M. Jeon, S. Kim, Y. Kwon and H. J. Kim (2020). “The role of TDP-43 propagation in neurodegenerative diseases: integrating insights from clinical and experimental studies.” Exp Mol Med 52(10): 1652–1662.

Johnson, M. L., A. A. Nagengast and H. K. Salz (2010). “PPS, a large multidomain protein, functions with sex-lethal to regulate alternative splicing in Drosophila.” PLoS Genet 6(3): e1000872.

Kaisinger, L. R., K. A. Kentistou, S. Stankovic, E. J. Gardner, F. R. Day, Y. Zhao, A. Mörseburg, C. J. Carnie, G. Zagnoli-Vieira, F. Puddu, S. P. Jackson, S. O’Rahilly, I. S. Farooqi, L. Dearden, L. C. Pantaleão, S. E. Ozanne, K. K. Ong and J. R. B. Perry (2023). “Large-scale exome sequence analysis identifies sex- and age-specific determinants of obesity.” Cell Genomics 3(8).

Kannan, K. and Y. W. Fridell (2013). “Functional implications of Drosophila insulin-like peptides in metabolism, aging, and dietary restriction.” Front Physiol 4: 288.

Kapan, N., O. V. Lushchak, J. Luo and D. R. Nässel (2012). “Identified peptidergic neurons in the Drosophila brain regulate insulin-producing cells, stress responses and metabolism by coexpressed short neuropeptide F and corazonin.” Cell Mol Life Sci 69(23): 4051–4066.

Kelly, S., C. Pak, M. Garshasbi, A. Kuss, A. H. Corbett and K. Moberg (2012). “New kid on the ID block: neural functions of the Nab2/ZC3H14 class of Cys₃His tandem zinc-finger polyadenosine RNA binding proteins.” RNA Biol 9(5): 555–562.

Kelly, S. M., R. Bienkowski, A. Banerjee, D. J. Melicharek, Z. A. Brewer, D. R. Marenda, A. H. Corbett and K. H. Moberg (2016). “The Drosophila ortholog of the Zc3h14 RNA binding protein acts within neurons to pattern axon projection in the developing brain.” Dev Neurobiol 76(1): 93–106.

Kelly, S. M., S. W. Leung, L. H. Apponi, A. M. Bramley, E. J. Tran, J. A. Chekanova, S. R. Wente and A. H. Corbett (2010). “Recognition of polyadenosine RNA by the zinc finger domain of nuclear poly(A) RNA-binding protein 2 (Nab2) is required for correct mRNA 3’-end formation.” J Biol Chem 285(34): 26022–26032.

Kelly, S. M., S. W. Leung, C. Pak, A. Banerjee, K. H. Moberg and A. H. Corbett (2014). “A conserved role for the zinc finger polyadenosine RNA binding protein, ZC3H14, in control of poly(A) tail length.” Rna 20(5): 681–688.

Kim, J. and T. P. Neufeld (2015). “Dietary sugar promotes systemic TOR activation in Drosophila through AKH-dependent selective secretion of Dilp3.” Nat Commun 6: 6846.

Kim, S. K., D. D. Tsao, G. S. B. Suh and I. Miguel-Aliaga (2021). “Discovering signaling mechanisms governing metabolism and metabolic diseases with Drosophila.” Cell Metab 33(7): 1279–1292.

Kishore, S., S. Luber and M. Zavolan (2010). “Deciphering the role of RNA-binding proteins in the post-transcriptional control of gene expression.” Brief Funct Genomics 9(5-6): 391–404.

Kréneisz, O., X. Chen, Y. W. Fridell and D. K. Mulkey (2010). “Glucose increases activity and Ca2+ in insulin-producing cells of adult Drosophila.” Neuroreport 21(17): 1116–1120.

Kühnlein, R. P. (2012). “Thematic review series: Lipid droplet synthesis and metabolism: from yeast to man. Lipid droplet-based storage fat metabolism in Drosophila.” J Lipid Res 53(8): 1430–1436.

Lancaster, C. L., P. S. Yalamanchili, J. N. Goldy, S. W. Leung, A. H. Corbett and K. H. Moberg (2024). “The RNA-binding protein Nab2 regulates levels of the RhoGEF Trio to govern axon and dendrite morphology.” Mol Biol Cell 35(8): ar109.

Leboucher, A., D. F. Pisani, L. Martinez-Gili, J. Chilloux, P. Bermudez-Martin, A. Van Dijck, T. Ganief, B. Macek, J. A. J. Becker, J. Le Merrer, R. F. Kooy, E. Z. Amri, E. W. Khandjian, M. E. Dumas and L. Davidovic (2019). “The translational regulator FMRP controls lipid and glucose metabolism in mice and humans.” Mol Metab 21: 22–35.

Lee, W. H., E. Corgiat, J. C. Rounds, Z. Shepherd, A. H. Corbett and K. H. Moberg (2020). “A Genetic Screen Links the Disease-Associated Nab2 RNA-Binding Protein to the Planar Cell Polarity Pathway in Drosophila melanogaster.” G3 (Bethesda) 10(10): 3575–3583.

Leung, S. W., L. H. Apponi, O. E. Cornejo, C. M. Kitchen, S. R. Valentini, G. K. Pavlath, C. M. Dunham and A. H. Corbett (2009). “Splice variants of the human ZC3H14 gene generate multiple isoforms of a zinc finger polyadenosine RNA binding protein.” Gene 439(1-2): 71–78.

Mauvais-Jarvis, F. (2015). “Sex differences in metabolic homeostasis, diabetes, and obesity.” Biol Sex Differ 6: 14.

Mircsof, D., M. Langouët, M. Rio, S. Moutton, K. Siquier-Pernet, C. Bole-Feysot, N. Cagnard, P. Nitschke, L. Gaspar, M. Žnidarič, O. Alibeu, A. K. Fritz, D. P. Wolfer, A. Schröter, G. Bosshard, M. Rudin, C. Koester, F. Crestani, P. Seebeck, N. Boddaert, K. Prescott, R. Hines, S. J. Moss, J. M. Fritschy, A. Munnich, J. Amiel, S. A. Brown, S. K. Tyagarajan and L. Colleaux (2015). “Mutations in NONO lead to syndromic intellectual disability and inhibitory synaptic defects.” Nat Neurosci 18(12): 1731–1736.

Moon, S. J., Y. Hu, M. Dzieciatkowska, A. R. Kim, J. M. Asara, A. D’Alessandro and N. Perrimon (2026). “Modeling tissue-specific Drosophila metabolism identifies high sugar diet-induced metabolic dysregulation in muscle at reaction and pathway levels.” Nat Commun 17(1): 1692.

Morton, G. J., D. E. Cummings, D. G. Baskin, G. S. Barsh and M. W. Schwartz (2006). “Central nervous system control of food intake and body weight.” Nature 443(7109): 289–295.

Nässel, D. R., O. I. Kubrak, Y. Liu, J. Luo and O. V. Lushchak (2013). “Factors that regulate insulin producing cells and their output in Drosophila.” Front Physiol 4: 252.

Oh, Y., J. S. Lai, H. J. Mills, H. Erdjument-Bromage, B. Giammarinaro, K. Saadipour, J. G. Wang, F. Abu, T. A. Neubert and G. S. B. Suh (2019). “A glucose-sensing neuron pair regulates insulin and glucagon in Drosophila.” Nature 574(7779): 559–564.

Pak, C., M. Garshasbi, K. Kahrizi, C. Gross, L. H. Apponi, J. J. Noto, S. M. Kelly, S. W. Leung, A. Tzschach, F. Behjati, S. S. Abedini, M. Mohseni, L. R. Jensen, H. Hu, B. Huang, S. N. Stahley, G. Liu, K. R. Williams, S. Burdick, Y. Feng, S. Sanyal, G. J. Bassell, H. H. Ropers, H. Najmabadi, A. H. Corbett, K. H. Moberg and A. W. Kuss (2011). “Mutation of the conserved polyadenosine RNA binding protein, ZC3H14/dNab2, impairs neural function in Drosophila and humans.” Proc Natl Acad Sci U S A 108(30): 12390–12395.

Penalva, L. O. and L. Sánchez (2003). “RNA binding protein sex-lethal (Sxl) and control of Drosophila sex determination and dosage compensation.” Microbiol Mol Biol Rev 67(3): 343–359, table of contents.

Post, S., G. Karashchuk, J. D. Wade, W. Sajid, P. De Meyts and M. Tatar (2018). “Drosophila Insulin-Like Peptides DILP2 and DILP5 Differentially Stimulate Cell Signaling and Glycogen Phosphorylase to Regulate Longevity.” Front Endocrinol (Lausanne) 9: 245.

Power, M. L. and J. Schulkin (2008). “Sex differences in fat storage, fat metabolism, and the health risks from obesity: possible evolutionary origins.” Br J Nutr 99(5): 931–940.

Rounds, J. C., E. B. Corgiat, C. Ye, J. A. Behnke, S. M. Kelly, A. H. Corbett and K. H. Moberg (2022). “The disease-associated proteins Drosophila Nab2 and Ataxin-2 interact with shared RNAs and coregulate neuronal morphology.” Genetics 220(1).

Schindelin, J., I. Arganda-Carreras, E. Frise, V. Kaynig, M. Longair, T. Pietzsch, S. Preibisch, C. Rueden, S. Saalfeld, B. Schmid, J.-Y. Tinevez, D. J. White, V. Hartenstein, K. Eliceiri, P. Tomancak and A. Cardona (2012). “Fiji: an open-source platform for biological-image analysis.” Nature Methods 9(7): 676–682.

Schmidt, U., M. Weigert, C. Broaddus and G. Myers (2018). Cell Detection with Star-Convex Polygons, Cham, Springer International Publishing.

Semaniuk, U., O. Strilbytska, K. Malinovska, K. B. Storey, A. Vaiserman, V. Lushchak and O. Lushchak (2021). “Factors that regulate expression patterns of insulin-like peptides and their association with physiological and metabolic traits in Drosophila.” Insect Biochem Mol Biol 135: 103609.

Shelby, G., A. H. Corbett and R. R. Parker (2026). “The role of ZC3H14 (MSUT2) in neurodevelopment and tauopathies.” Brain.

Soucek, S., Y. Zeng, D. L. Bellur, M. Bergkessel, K. J. Morris, Q. Deng, D. Duong, N. T. Seyfried, C. Guthrie, J. P. Staley, M. B. Fasken and A. H. Corbett (2016). “The Evolutionarily-conserved Polyadenosine RNA Binding Protein, Nab2, Cooperates with Splicing Machinery to Regulate the Fate of pre-mRNA.” Mol Cell Biol 36(21): 2697–2714.

Sudhakar, S. R., H. Pathak, N. Rehman, J. Fernandes, S. Vishnu and J. Varghese (2020). “Insulin signalling elicits hunger-induced feeding in Drosophila.” Dev Biol 459(2): 87–99.

Teleman, A. A. (2009). “Molecular mechanisms of metabolic regulation by insulin in Drosophila.” Biochem J 425(1): 13–26.

Uchewa, O. O., E. J. Alobu, F. C. Ikechukwu, J. C. Udoadi, C. C. Jachike, M. C. Ifeanyi, F. C. Ibenne, C. P. Iheme, P. N. Ihedi, C. Item, C. D. Isaiah, S. E. Irem and A. O. Ibegbu (2025). “The role of glia cell in the neural mechanism of memory formation, storage, and motor control: A review.” Neuroscience 588: 174–192.

Varlamov, O., C. L. Bethea and C. T. Roberts, Jr. (2014). “Sex-specific differences in lipid and glucose metabolism.” Front Endocrinol (Lausanne) 5: 241.

Viola, C. M., O. Frittmann, H. T. Jenkins, T. Shafi, P. De Meyts and A. M. Brzozowski (2023). “Structural conservation of insulin/IGF signalling axis at the insulin receptors level in Drosophila and humans.” Nat Commun 14(1): 6271.

Walls, S. M., Jr., S. J. Attle, G. B. Brulte, M. L. Walls, K. D. Finley, D. A. Chatfield, D. R. Herr and G. L. Harris (2013). “Identification of sphingolipid metabolites that induce obesity via misregulation of appetite, caloric intake and fat storage in Drosophila.” PLoS Genet 9(12): e1003970.

Wat, L. W., C. Chao, R. Bartlett, J. L. Buchanan, J. W. Millington, H. J. Chih, Z. S. Chowdhury, P. Biswas, V. Huang, L. J. Shin, L. C. Wang, M. L. Gauthier, M. C. Barone, K. L. Montooth, M. A. Welte and E. J. Rideout (2020). “A role for triglyceride lipase brummer in the regulation of sex differences in Drosophila fat storage and breakdown.” PLoS Biol 18(1): e3000595.

Wheeler, J. M., P. McMillan, T. J. Strovas, N. F. Liachko, A. Amlie-Wolf, R. L. Kow, R. L. Klein, P. Szot, L. Robinson, C. Guthrie, A. Saxton, N. M. Kanaan, M. Raskind, E. Peskind, J. Q. Trojanowski, V. M. Y. Lee, L. S. Wang, C. D. Keene, T. Bird, G. D. Schellenberg and B. Kraemer (2019). “Activity of the poly(A) binding protein MSUT2 determines susceptibility to pathological tau in the mammalian brain.” Sci Transl Med 11(523).

Xiao, F. and F. Guo (2022). “Impacts of essential amino acids on energy balance.” Mol Metab 57: 101393.

Yamaguchi, T., R. Fernandez and R. A. Roth (1995). “Comparison of the signaling abilities of the Drosophila and human insulin receptors in mammalian cells.” Biochemistry 34(15): 4962–4968.

Zhou, W., L. Zhao, Z. Mao, Z. Wang, Z. Zhang and M. Li (2023). “Bidirectional Communication Between the Brain and Other Organs: The Role of Extracellular Vesicles.” Cell Mol Neurobiol 43(6): 2675–2696.

Zou, J., J. Li, X. Wang, D. Tang and R. Chen (2024). “Neuroimmune modulation in liver pathophysiology.” J Neuroinflammation 21(1): 188.

